# The MTL200: a surface-based, probabilistic atlas of the medial temporal lobe

**DOI:** 10.64898/2025.12.04.692420

**Authors:** Ian Faul, Adya Agarwal, Stephen Graziose, Andrew Nwacha, Janis Park, Anisah Sahibul, Katie Tong, Ben Deen

## Abstract

The human medial temporal lobe (MTL) includes a number of cortical regions surrounding the hippocampus implicated in long-term memory and high-level perception, including temporopolar cortex (TPC), entorhinal cortex (ERC), perirhinal cortex (PRC), and parahippocampal cortex (PHC). A critical prerequisite to studying the function of these areas is to define their anatomical boundaries. While several atlases or segmentations methods for automated boundary definition exist, they have limitations: 1) they are derived from small sample sizes; 2) most do not include TPC; and 3) they are volumetric rather than surface-based and are therefore not straightforward to use for surface-based analysis of MRI data. Here we present the MTL200: a surface-based, probabilistic atlas of MTL regions TPC, ERC, PRC, and PHC. The atlas is derived from hand drawings of regions based on gross anatomical features corresponding to cytoarchitectonic boundaries, using anatomical MRI data from 200 human participants. This work provides a valuable tool for researchers studying the anatomy and function of MTL regions, particularly the poorly characterized area TPC.

## 1. Introduction

The primate medial temporal lobe (MTL) comprises a set of interconnected brain areas considered essential for declarative long-term memory (Cohen & Eichenbaum, 1993; Squire & Zola-Morgan, 1991). These include the hippocampal formation and adjacent areas – entorhinal (ERC), perirhinal (PRC), parahippocampal cortex (PHC), and temporopolar cortex (TPC) – that provide an interface between the hippocampus and neocortex (Suzuki, 1996). In order to study the function of these areas in humans, their anatomical location must be determined. The current report describes a novel, surface-based, probabilistic atlas that can be used to facilitate the identification of these areas from structural MRI data.

Subregions of the MTL are classically defined in terms of cytoarchitecture, connectivity, and function. Our understanding of the functional organization of the primate MTL has been heavily informed by studies of anatomical connectivity in the macaque. ERC provides the dominant cortical input to the hippocampal formation via the perforant pathway. ERC receives a majority of input through bidirectional connections with PRC, PHC, and TPC (Insausti et al., 1987; Suzuki & Amaral, 1994). In turn, PRC, PHC, and TPC are connected to a distinct set of cortical regions. PRC, or Brodmann’s area (BA) 35/36, is primarily connected to areas within the ventral visual stream (Suzuki & Amaral, 1994; Van Hoesen & Pandya, 1975) and is thought to carry high-level sensory information such as object identity (Buffalo et al., 1998; Murray & Richmond, 2001). PHC, or area TH/TF, has a widely distributed set of connections to frontal, parietal, and temporal areas (Seltzer & Pandya, 1976; Suzuki & Amaral, 1994), with a spatial organization resembling the default mode network (Binder et al., 2009; Buckner & Margulies, 2019). PHC has been argued to carry information about spatial and contextual relationships (Aminoff et al., 2013; Eichenbaum & Lipton, 2008; Epstein, 2008). Prominent theoretical models of MTL function argue that distinct streams of information relayed through PRC and PHC are integrated by the hippocampus, supporting the formation of relational long-term memories (Davachi, 2006; Eichenbaum et al., 2007; Ranganath & Ritchey, 2012).

TPC, also termed BA38 or TG, has figured less prominently into theories of hippocampal-dependent memory and isn’t consistently included in definitions of the MTL. However, the connectivity and function of this region indicate that it should be considered along with PRC and PHC as an interface between the hippocampus and neocortex. TPC in the macaque has a direct, bidirectional connection with entorhinal cortex, similar in magnitude to the projections from PRC and PHC (Insausti et al., 1987; Moran et al., 1987). Some studies in the macaque referred to TPC as a subregion of PRC termed area 36 polar (Amaral et al., 1987; Insausti et al., 1987). Functionally, TPC has been implicated in long-term memory for social and semantic information: for instance, humans with TPC dysfunction resulting from brain damage or neural degeneration can have deficits naming familiar objects or faces (Barton et al., 2001; Damasio et al., 1990; Gentileschi et al., 2001). While TPC has been difficult to study using human neuroimaging due to signal artifacts in fMRI data from this area (Olson et al., 2007), recent work has used optimized data acquisition approaches to improve signal quality, enabling further study of TPC’s function and contribution to memory (Deen et al., 2024; Girn et al., 2024).

In order to study the function of MTL regions in humans, a key prerequisite is to delineate their anatomical boundaries. To this end, prior work has developed protocols for defining the boundaries for TPC, ERC, PRC, and PHC based on gross anatomical landmarks that correspond to cytoarchitectonic boundaries established in postmortem specimens (Augustinack et al., 2013; Feczko et al., 2009; Goncharova et al., 2001; Insausti et al., 1998; Pruessner et al., 2002). These protocols allow boundaries of MTL regions to be drawn on structural MRI data acquired noninvasively in vivo. However, hand-drawing areal boundaries is a slow, laborious process that cannot realistically be applied to the large-scale datasets that are increasingly common in MRI research, involving 1,000 or more participants (Bycroft et al., 2018; Casey et al., 2018; Jack Jr et al., 2008; Van Essen et al., 2013).

As an alternative, atlases derived from separate groups of participants can be used to infer the likely location of MTL subregions in a new MRI dataset. Several atlases or automated segmentation protocols for MTL subregions have been developed (Bouyeure et al., 2018; Iglesias et al., 2015; Ravikumar et al., 2021; Wisse et al., 2016; Xie et al., 2016; Yushkevich, Pluta, et al., 2015). The current work describes a new, publicly available atlas of neocortical areas within the MTL that complements existing options in three ways: the atlas is 1) surface-based; 2) derived from a large sample (*N* = 200), and 3) includes TPC.

Existing atlases of the MTL based on cytoarchitectonic boundaries are formatted as 3D volumetric datasets (Bouyeure et al., 2018; Yushkevich, Pluta, et al., 2015). However, functional MRI data are often analyzed after resampling to a 2D cortical surface representation (Dale et al., 1999; Fischl, Sereno, & Dale, 1999). Surface-based analysis of fMRI data uses a coordinate system that aligns with the natural geometry of cortex, and offers numerous technical advantages over volume-based analysis: improved intersubject alignment, the ability to spatially smooth across the surface, improved sensitivity to activation, and more efficient data storage (Anticevic et al., 2008; Coalson et al., 2018; Jo et al., 2007; Klein et al., 2010). The use of surface-based analysis has grown in the past decade through the impact of the Human Connectome Project (HCP), which developed an optimized analysis approach along with software for its implementation (Glasser, Smith, et al., 2016). While existing discrete parcellations of the cortical surface include areas in the MTL (Desikan et al., 2006; Destrieux et al., 2010; Fischl et al., 2004; Glasser, Coalson, et al., 2016), these areas are not defined based on cytoarchitectonic boundaries, and it remains unclear how they correspond with cytoarchitectonically defined areas from volumetric atlases (Insausti et al., 1998; Yushkevich, Pluta, et al., 2015). The present work introduces an atlas of the MTL that is natively formatted in surface-based coordinate spaces fsaverage and fsLR, and can be easily applied to analyze MRI data that has been registered to these spaces using Freesurfer or the HCP pipelines.

A second limitation of existing MTL atlases is the relatively small sample size they are derived from. Current atlases have been built from samples in the range of 15 – 40 human participants (Amunts et al., 2020; Bouyeure et al., 2018; Yushkevich, Pluta, et al., 2015). While justified by the substantial amount of time required to hand-label anatomical boundaries, small sample sizes limit the ability to evaluate the uncertainty of areal boundaries given variation across participants in neuroanatomy and registration quality. In particular, the collateral sulcus, which factors into the boundaries of all neocortical MTL regions, exhibits substantial variation across individuals in its length, branching, and gaps (Feczko et al., 2009; Pruessner et al., 2002). In order to robustly sample the range of anatomical variation present in healthy adults, the current work uses a large set of 200 participants.

Lastly, most existing MTL atlases based on cytoarchitectonic boundaries include ERC, PRC, and PHC, but not TPC (Ravikumar et al., 2021; Wisse et al., 2016; Xie et al., 2016; Yushkevich, Pluta, et al., 2015), with the exception of one atlas derived from a pediatric sample (Bouyeure et al., 2018). Here, we include TPC in our parcellation, aiming to facilitate further study of this poorly understood area.

To develop the MTL200 atlas, MTL regions were hand-drawn on volumetric anatomical images of 200 participants from the HCP dataset, based on gross anatomical landmarks previously established to capture cytoarchitectonic boundaries (Insausti et al., 1998). They were then resampled to a 2D cortical surface reconstruction for each participant and registered to surface-based atlases fsLR and fsaverage. Lastly, regions in surface space were combined across participants to generate probability maps for each region, as well as a discrete segmentation based on maximum likelihood. The resulting atlas is publicly available (https://github.com/SocialMemoryLab/MTL200) and will provide a resource for researchers studying the structure and function of the MTL using surface-based analysis of MRI data.

## 2. Methods

### 2.1 Participants

To generate the atlas, we used data from *N* = 200 unrelated young adult participants from the HCP S1200 release (https://www.humanconnectome.org/study/hcp-young-adult/document/1200-subjects-data-release). To select a subset of participants, we used the following criteria: 1) participants missing anatomical data were excluded; 2) participants with reported quality control issues were excluded; 3) an equal number of males and females were chosen; and 4) the age distributions for males and females were matched as closely as possible. The resulting sample comprised 100 males and 100 females with an average age of 29 yrs (Table 1). All participants were right-handed and had no history of medical condition. Participants provided written, informed consent, and the protocol was approved by the Washington University institutional review board.

**Table 1:**
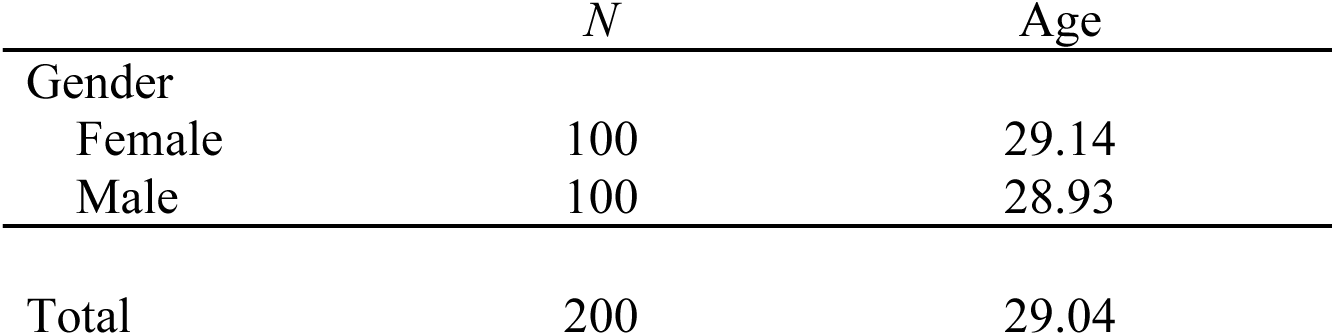
Sample demographics.

|  | *N* | Age |
| --- | --- | --- |
| Gender |  |  |
| Female | 100 | 29.14 |
| Male | 100 | 28.93 |
| Total | 200 | 29.04 |

### 2.2 MRI data acquisition

MRI data were acquired on a Siemens 3T Skyra scanner, using HCP’s data acquisition protocols (Van Essen et al., 2012). Structural MRI data were used to hand-draw anatomical regions, and resting-state functional data were used to assess functional connectivity for the sake of atlas validation. T1-weighted anatomical images were acquired using a 3D magnetization-prepared rapid acquisition gradient echo (*MPRAGE*) sequence with inversion recovery, with echo time (TE) = 2.14ms, repetition time (TR) = 2400ms, inversion time = 1000ms, flip angle (FA) = 8°, bandwidth (BW) = 210Hz/pixel, echo spacing (ES) = 7.6ms, field of view (FOV) = 180 × 224 × 224mm^3^, and voxel size = 0.7mm isotropic. T2-weighted anatomical images were acquired using a 3D SPACE sequence with TE = 565ms, TR = 3200ms, variable FA, FOV = 180 x 224 x 224 mm^3^, and voxel size = 0.7mm isotropic. fMRI data were acquired using a gradient echo echo-planar-imaging sequence, with TE = 33.1ms, TR = 720ms, FA = 52°, BW = 2290Hz/pixel, ES = 0.58ms, FOV = 208×180mm^2^, voxel size = 2mm isotropic, and a multiband acceleration factor of 8. Resting-state data were collected from 197 of the 200 participants. For most participants, this data included four runs with 1,200 time points each. 1 participant had only 3 runs; 12 participants had 2 runs; and 1 participant had 1 run.

### 2.3 MRI data preprocessing

Structural and functional MRI data were preprocessed using the HCP’s minimal preprocessing pipeline (Glasser et al., 2013). A summary will be given here, with further details provided in prior publication. Preprocessing relied on tools from the FMRIB Software Library (FSL) and Freesurfer (Fischl, 2012; Smith et al., 2004). Images were first corrected for distortions induced by gradient nonlinearities. Repeated acquisition of anatomical images were registered and averaged. The resulting anatomical images were rigidly registered (6-DOF transformation) to the MNI152 template image used by FSL (MNI152NLin6Asym), and bias corrected. Freesurfer’s recon-all was used to generated a triangle mesh reconstruction of the cortical surface (Dale et al., 1999; Fischl, Sereno, & Dale, 1999). Surface coordinates were registered to the fsaverage and fsLR template spaces (Fischl, Sereno, Tootell, et al., 1999), with Freesurfer’s initial registration further optimized using multimodal surface matching (Robinson et al., 2014). A nonlinear volumetric registration to MNI152 space was also computed using FNIRT.

For anatomical images, one additional step was performed beyond the HCP’s preprocessing. Studies of cytoarchitectonic regions within the MTL have typically characterized these areas on coronal slices aligned perpendicularly to the axis of the hippocampus and temporal lobe (Insausti et al., 1998). Protocols for identifying subregions using gross anatomical features with validation from cytoarchitectonic boundaries identified ex-vivo have used this brain orientation. However, the orientation of the brain in MNI space differs, with the hippocampus angled obliquely in the *y*-*z* plane. To account for this difference, we applied a counterclockwise rotation of 43.3° about the x-axis to all anatomical images. This specific angle was chosen based on the MNI152 template brain as the angle that brought the hippocampus into alignment with the *y*-axis. The resulting coordinate system will be referred to as hippocampus (HC)-aligned space.

fMRI data were first corrected for distortions induced by gradient nonlinearity. Parameters for motion correction were estimated using rigid transformations from FSL’s MCFLIRT (Jenkinson et al., 2002). A transformation to correct EPI distortion was estimated using FSL’s topup (Andersson et al., 2003). Functional to anatomical registration was performed using Freesurfer’s bbregister (Greve & Fischl, 2009). Transformations applying motion correction, distortion correction, and registration were combined and applied in a single resampling step. Noise signal was removed from data using independent component analysis (ICA-FIX) as implemented in the HCP pipeline (Salimi-Khorshidi et al., 2014). fMRI data were resampled from volumetric to surface space using ribbon-constrained mapping, and then resampled to the fsLR atlas at 32k vertex density (Glasser et al., 2013).

### 2.4 Manual labeling

MTL subregions TPC, ERC, PRC, and PHC bilaterally were hand-drawn on each participant’s HC-aligned anatomical image by five trained researchers (authors I.F., S.G., A.N., J.P., and K.T.). Participants were pseudorandomly assigned to each researcher. Drawings for 28 participants were completed by I.F., 52 by S.G., 31 by A.N., 33 by J.P., and 56 by K.T.

We established a labeling procedure that enabled the efficient generation of labels for a large number of participants, while taking advantage of the accurate surface reconstructions provided by the HCP pipeline. Manually labeling regions at the native anatomical resolution of .7mm would be impractical with a large sample and would involve hand-tracing gray-white and gray-pial boundaries that were already established by the preprocessing pipeline. Instead, regions were drawn on a 2mm-resolution grid overlaid on .7mm-resolution anatomical images. Researchers were instructed to be inclusive of gray matter even if the drawn regions included white matter or CSF, since components outside of gray matter would be removed through surface resampling. Lastly, to generate accurate volumetric masks for producing a volumetric atlas and measuring areal volumes, regions were resampled from the surface back to the volume (detailed below in Atlas Construction section).

Boundaries for each region were determined using gross anatomical features following the protocol of Insausti et al. (1998), as detailed below. The location of the collateral sulcus (CS), which played a key role in defining areal boundaries, was determined by visualizing both 3D volumetric images using fsleyes, and surface reconstructions using the Connectome Workbench tools. Regions were hand-drawn using fsleyes, producing binary mask files with value 1 inside the region and 0 outside.

#### 2.4.1 Parahippocampal cortex

The posterior boundary of PHC was defined as the last coronal slice in which the HC could be observed. The anterior boundary was defined as 4mm (2 slices) posterior to the posterior-most slice containing the head of the HC, i.e. the boundary between the hippocampal head and body. The medial boundary of PHC was defined as the medial-most point of the parahippocampal gyrus (PHG), and the lateral boundary was the lateral edge of the CS.

#### 2.4.2 Perirhinal and entorhinal cortices

The posterior boundary of PRC was defined as the first slice anterior to the PHC. PRC surrounds ERC in the anterior, lateral, and posterior directions, extending 2mm beyond ERC in the anterior and posterior directions. Therefore the first and last 2mm slice within the anterior-posterior extent of PRC/ERC was defined as PRC. All slices in between contained both PRC (laterally) and ERC (medially). The medial boundary of PRC/ERC was defined as the medial-most point of the PHG, and the lateral boundary as the lateral edge of the CS. In slices containing ERC, the boundary between PRC and ERC was positioned within the CS, with its precise location determined by the depth of the CS, following the convention of Insausti et al. (1998). For a CS with typical depth (1-1.5cm), the lateral border of ERC was defined as the midpoint of the medial bank of the CS. For a shallow CS (<1cm), the lateral border of ERC was defined as the fundus of the CS. For a deep CS (>1.5mm) the lateral border of ERC was defined as the medial edge of the CS. We note that other protocols for MRI-based delineation of ERC and PRC have used alternative definitions for this boundary that don’t depend on CS depth, such as the fundus of the CS (Honeycutt et al., 1998) or the medial edge of the CS (Augustinack et al., 2013; Berron et al., 2017; Feczko et al., 2009; Goncharova et al., 2001).

#### 2.4.3 Temporopolar cortex

TPC lies at the anterior-most extent of the temporal lobe. The term TPC has been used variably in the literature, to denote either a cytoarchitectonically defined area restricted largely to the medial surface of the temporal lobe (Blaizot et al., 2010; Insausti et al., 1998), or a broader region encompassing multiple cytoarchitectonic areas on the anterior medial and lateral surfaces (Ding et al., 2009). The current atlas follows the convention of Insausti et al. (1998). The posterior boundary of TPC was defined as the farther anterior of the following two landmarks: 1) the anterior-most point of the CS; 2) 2mm (one slice) anterior to the fronto-temporal junction (FTJ), where the FTJ is defined as the anterior-most slice in which white matter connecting the frontal and temporal lobes is present. In almost all participants in our sample, the TPC/PRC boundary was defined by the first landmark. At the anterior-most part of TPC (typically 6-8mm), all gray matter within the temporal lobe belongs to TPC. Posteriorly, TPC is restricted to the ventromedial surface. The dorsolateral boundary of TPC was defined by the fundus of the lateral-most polar sulcus, i.e. the sulcus after the last gyrus of Schwalbe. The ventrolateral boundary of TPC was defined as the medial edge of the sulcus just lateral to the CS (typically the inferior temporal sulcus [ITS], but occasionally the superior temporal sulcus [STS]).

### 2.5 Atlas construction

Hand-drawn regions were next resampled to the cortical surface. Interpolation was performed using ribbon-constrained mapping, an algorithm developed as part of the HCP that uses white and pial surfaces to determine which voxels to sample from (Glasser et al., 2013). Areas were first resampled to each participant’s native surface produced by Freesurfer, and then transformed to fsLR space using barycentric resampling. Topological defects, such as holes and islands, that were produced in the volume-to-surface resampling step were corrected using the Freesurfer functions -metric-fill-holes and -remove-islands in each subject. Lastly, mask files were averaged across participants to produce a probability map for each region, specifying the likelihood of the region existing at a given coordinate. These maps were also resampled to fsaverage to provide the atlas in multiple template spaces.

In addition to probability maps for each region, we generated a discrete segmentation corresponding to a maximum likelihood estimate of which of the four regions exists at a given coordinate. To generate the discrete segmentation, we first generated a mask covering the entire MTL. For each participant and hemisphere, mask files specifying the four areas on the surface were combined to generate an MTL mask. These MTL masks were averaged across participants to generate a probability map. To generate a group-level MTL mask for each hemisphere, we thresholded this probability map at a value chosen so that the size (number of vertices) of the resulting mask was equal to the mean size of MTL masks across participants. Within the group-level MTL mask, each coordinate was assigned to one of the four regions based on maximum likelihood, resulting in a discrete segmentation.

While the focus of this manuscript is on generating a surface-based atlas, we additionally generated a volumetric atlas in MNI152 space that can be used by researchers conducting volumetric MRI analysis. Within each participant, hand-drawn masks were resampled from the surface to the .7mm anatomical image using ribbon-constrained mapping, generating a volume-based mask at higher resolution than the original hand drawings. Volumetric masks were transformed to MNI152 space using the nonlinear transformation generated by the HCP pipeline. Lastly, masks were averaged across participants to produce probability maps.

### 2.6 Data analysis

#### 2.6.1 Reliability measures

Two procedures were used to assess the reliability of hand-drawn regions. First, inter-rater reliability was measured in a subset of *N* = 35 participants for whom regions were drawn by two researchers (I.F. and K.T.). Surface-based masks were compared across the two researchers using the Dice coefficient over surface vertices.

Second, we conducted a split-half reliability analysis to determine the consistency of probability maps and discrete segmentations across subsets of participants. We partitioned participants into two halves with matched age, sex, and rater distributions. Surface-based probability maps and discrete segmentations were computed for each half. Reliability of probability maps was assessed by computing correlations among non-zero coordinates in each map. Reliability of regions in discrete segmentations was assessed using the Dice coefficient.

#### 2.6.2 Comparison of regional volumes and surface extent

Gray matter volumes for MTL subregions were computed from 0.7mm volumetric masks. The mean volumes were compared with prior studies using similar protocols (Insausti et al., 1998; Pruessner et al., 2002). A paired, two-sample *t*-test was used to evaluate hemispheric differences in volume for each region. Bonferroni correction was used to account for multiple comparisons, applying a significance level of α = .05/4 = .0125. The size and extent of MTL subregions on the cortical surface was compared with existing discrete, surface-based segmentations that included the MTL (Desikan et al., 2006; Fischl et al., 2004; Glasser, Coalson, et al., 2016).

#### 2.6.3 Functional connectivity

To provide an example of how the atlas can be used, and evaluate functional dissociations between MTL regions, we last measured resting-state functional connectivity of labeled regions. We used two complementary approaches: 1) a “classical” approach assessing group-averaged whole-brain functional connectivity maps; and 2) a modern approach using precision functional mapping, assessing connectivity to whole-brain functional networks defined in individual participants. These analyses included a subset of *N* = 197 participants for whom resting-state data was available.

For the first approach, seed regions were defined using the group-level discrete segmentation. For each participant, correlations were computed between time series from the seed region and each surface coordinate. Functional connectivity maps were then averaged across participants.

For the second approach, individualized functional network parcellations were first generated using the multi-session hierarchical Bayesian model (MSHBM) algorithm (Kong et al., 2019), using a 15-network parcellation as a group-level prior (Du et al., 2024). Time series correlations where then computed between MTL regions defined in individuals and the mean series from each functional network. A linear mixed effects model was used to statistically evaluate differences in connectivity profiles across MTL regions, with participant included as a random effect.

## 3. Results

We generated the MTL200: a surface-based, probabilistic atlas of MTL regions TPC, ERC, PRC, and PHC, by hand-labeling regions based on gross anatomical features. Probability maps for each region in fsLR surface space, showing the proportion of participants with a labeled region at each coordinate, are shown in Fig. 2a. A discrete map showing which region had the highest likelihood at a given coordinate is shown in Fig. 2b. Volumetric probability maps in MNI152 space are shown in Fig. S4.

**Figure 1:**
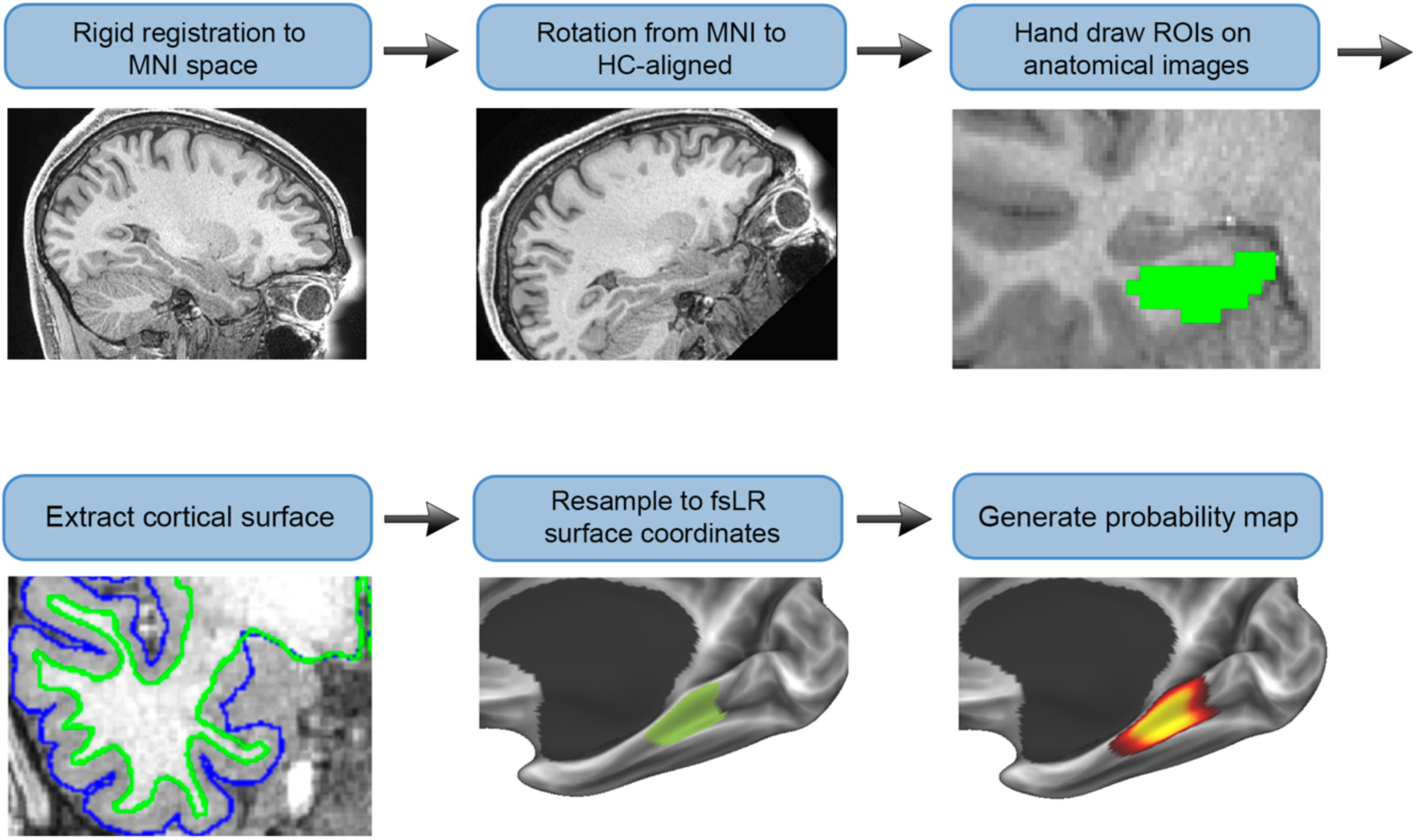
Atlas construction approach. Anatomical images were first rigidly aligned with MNI152 space, and rotated to align the hippocampus (HC) with the y-axis. Next, regions were hand-drawn on coronal slices. Cortical surface reconstructions were generated using Freesurfer. Regions were resampled from the volume to the surface (fsLR space). Lastly, regions were averaged across participants to produce a probability map.

**Figure 2:**
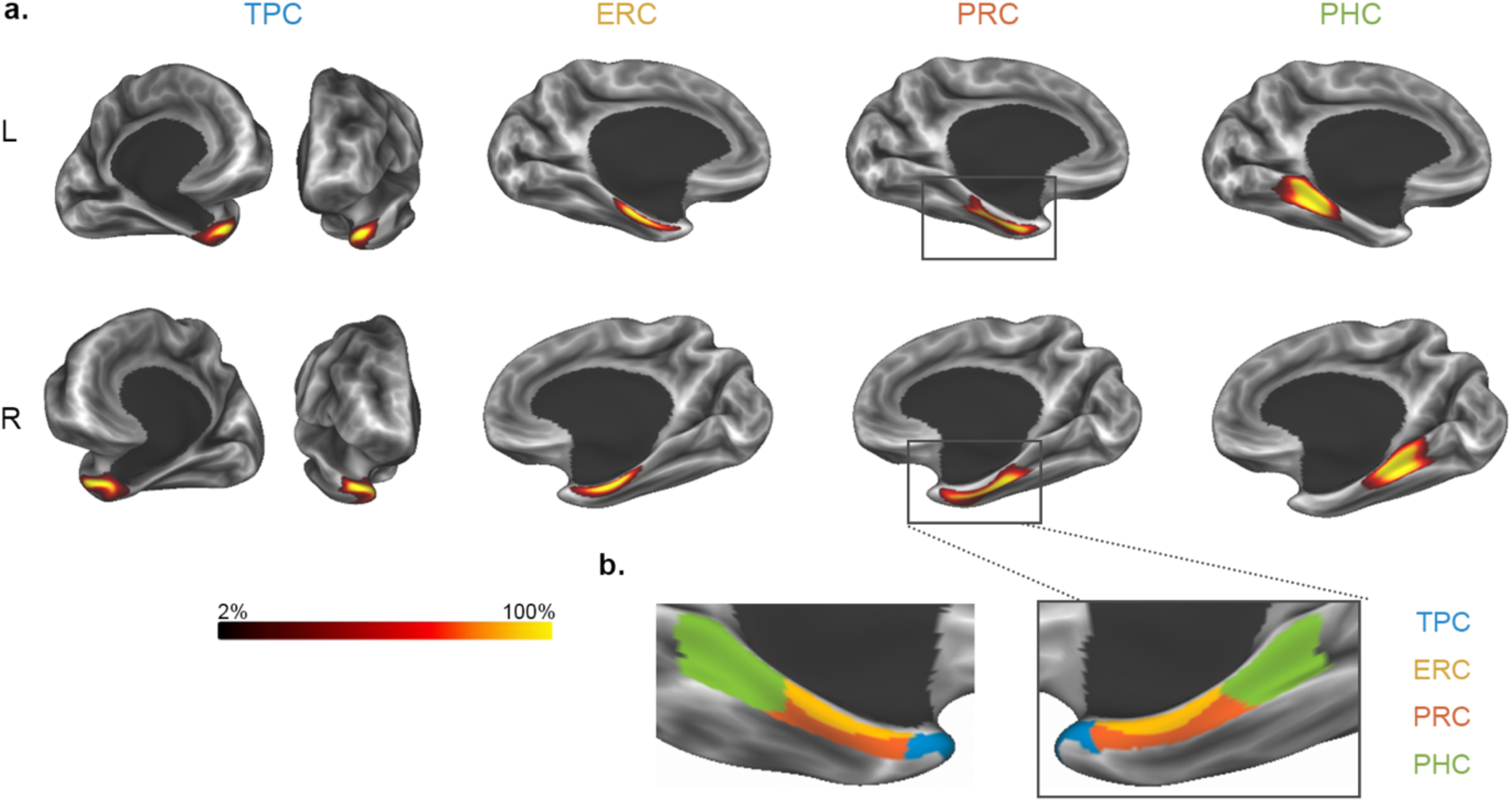
The MTL200 atlas. **a.** Probability maps for each region displayed on an inflated cortical surface. Regions include temporopolar cortex (TPC), entorhinal cortex (ERC), perirhinal cortex (PRC), and parahippocampal cortex (PHC). **b.** Discrete segmentations showing the most probable area for a given coordinate.

### 3.1 Interrater reliability

Inter-rater reliability was measured by having two researchers independently parcellate the same subset of 35 participants. Table 2 lists the average similarity scores (Dice coefficients) for each of the four MTL regions in each hemisphere. Interrater reliability values were high, ranging from .69 to .86 across regions and hemispheres (Table 2). This indicates that different researchers were able to consistently implement the protocol for identifying areal boundaries.

**Table 2.** Inter-rater reliability by region.

| Hemisphere | Region | Dice score |
| --- | --- | --- |
| L | TPC | 0.692 |
|  | ERC | 0.792 |
|  | PRC | 0.753 |
|  | PHC | 0.857 |
| R | TPC | 0.780 |
|  | ERC | 0.784 |
|  | PRC | 0.776 |
|  | PHC | 0.849 |

### 3.2 Split-half reliability

To further assess the reliability of our parcellation approach, we measured the similarity between atlases generated from two partitions of *N* = 100 participants matched their distribution of age, sex, and rater. Probabilistic atlases and discrete segmentations were generated using the same procedure as for the full dataset, as described in the Methods. Results were highly similar across the two partitions (Table 3). Dice coefficients for group-level discrete segmentations ranged from .94 to .98 across regions and hemispheres. Correlations between nonzero values in probability maps ranged from .96 to .99. These results demonstrate that our sample size is sufficiently large to generate highly reliable spatial profiles. This reliability is observed for both the most likely area at a given coordinate (discrete segmentation) and for continuous maps that incorporate variation in areal boundaries across participants (probability maps).

**Table 3.** Split-half reliability scores by region.

| Hemisphere | Region | Dice score | Correlation (r) |
| --- | --- | --- | --- |
| L | TPC | 0.935 | 0.964 |
|  | ERC | 0.972 | 0.970 |
|  | PRC | 0.952 | 0.965 |
|  | PHC | 0.975 | 0.973 |
| R | TPC | 0.947 | 0.969 |
|  | ERC | 0.947 | 0.977 |
|  | PRC | 0.948 | 0.970 |
|  | TPC | 0.947 | 0.969 |

### 3.3 Analysis of regional volumes

Volumes of labeled cortical regions were measured after resampling regions from the cortical surface back to each participant’s volumetric anatomical image (Table 4). Observed volumes were generally similar to prior reports (Insausti et al., 1998; Pruessner et al., 2002). TPC had a mean volume of 2587mm^3^ (left) and 2879 mm^3^ (right). These values are somewhat lower than previously reported mean TPC volumes from Insausti et al. (3421mm^3^ left, 3228mm^3^ right), but within the range of reported values. The values diverge from previously reported mean TPC volumes from Pruessner et al. (6224mm^3^ left, 6913mm^3^ right). ERC had a mean volume of 1277mm^3^ (left) and 1212mm^3^ (right). These values are somewhat lower than previously reported mean ERC volumes from Insausti et al. (1510 mm^3^ left, 1694mm^3^ right) and Pruessner et al. (1553mm^3^ left, 1672mm^3^ right), but within the range of values reported in both studies. PRC had a mean volume of 1801mm^3^ (left) and 1759mm^3^ (right). These values are somewhat lower than previously reported mean PRC volumes from Insausti et al. (2585mm^3^ left, 2577mm^3^ right) and Pruessner et al. (2502mm^3^ left, 2417mm^3^ right), but within the range of values reported in both studies. PHC had a mean volume of 2555mm^3^ (left) and 2450mm^3^ (right). These values are consistent with previously reported mean PHC volumes from Pruessner et al. (2675mm^3^ left, 2469mm^3^ right).

**Table 4.** Volumes of MTL subregions in mm^3^.

| Hemisphere | Region | Mean volume | Standard deviation |
| --- | --- | --- | --- |
| L | TPC | 2587 | 1078 |
|  | ERC | 1277 | 422 |
|  | PRC | 1801 | 527 |
|  | PHC | 2555 | 487 |
| R | TPC | 2879 | 1107 |
|  | ERC | 1212 | 405 |
|  | PRC | 1759 | 499 |
|  | PHC | 2450 | 479 |

Do the volumes of MTL regions differ across hemispheres? Our large sample of 200 participants is well suited to address this question. TPC was significantly larger in the right hemisphere (*t*[199] = −4.38, *P* < 0.0001), as was ERC (*t*[199] = 3.05, *P* < .005). PHC was significantly larger in the left hemisphere (*t*[199] = 3.39, *P* < .001). PRC did not show a significant difference in size across hemispheres (*t*[199] = 1.51, p = 0.1323).

### 3.4 Comparison with prior parcellations

Commonly used surface-based cortical parcellations, including the Glasser multimodal parcellation (MMP, Glasser, Coalson, et al., 2016), Desikan-Killiany atlas (DK, Desikan et al., 2006), and Destrieux atlas (DT, Destrieux et al., 2010), have substantial differences in the extent and location of labeled MTL regions. For instance, TPC ranges in size from 154 surface vertices in the DK atlas (left “temporalpole”) to 418 vertices in the MMP atlas (bilateral “TGd”; both regions assessed in 32k fsLR space). These atlases define areal boundaries using distinct criteria—either gross anatomical features (Desikan et al., 2006; Destrieux et al., 2010) or functional and anatomical connectivity profiles (Glasser, Coalson, et al., 2016)—but do not specifically use features related to cytoarchitectonic boundaries. How do these existing parcellations relate to the areal boundaries of the MTL200?

To compare parcellations, we visualized regions from the MTL200, MMP, DK, and DT on the cortical surface (Fig. 3). The extent of TPC varied dramatically across parcellations. As expected from the labeling criteria applied here, TPC in the MTL200 was largely restricted to the anterior and medial surfaces of the temporal lobe, and did not include the ITS, STS, or middle temporal gyrus (MTG). By contrast, the TGd region from MMP extended onto the lateral surface, including anterior components of lateral temporal structures ITS, STS, and MTG. The “temporalpole” region from DK was similar to MTL200, with minimal coverage of lateral temporal cortex. The “Pole_temporal” region from DT was intermediate in this regard, extending somewhat into lateral temporal cortex, but less so than the MMP’s TGd. These results demonstrate that some prior surface-based cortical parcellations include a TPC label that extends farther into lateral temporal cortex than would be expected from cytoarchitectonic boundaries.

**Figure 3:**
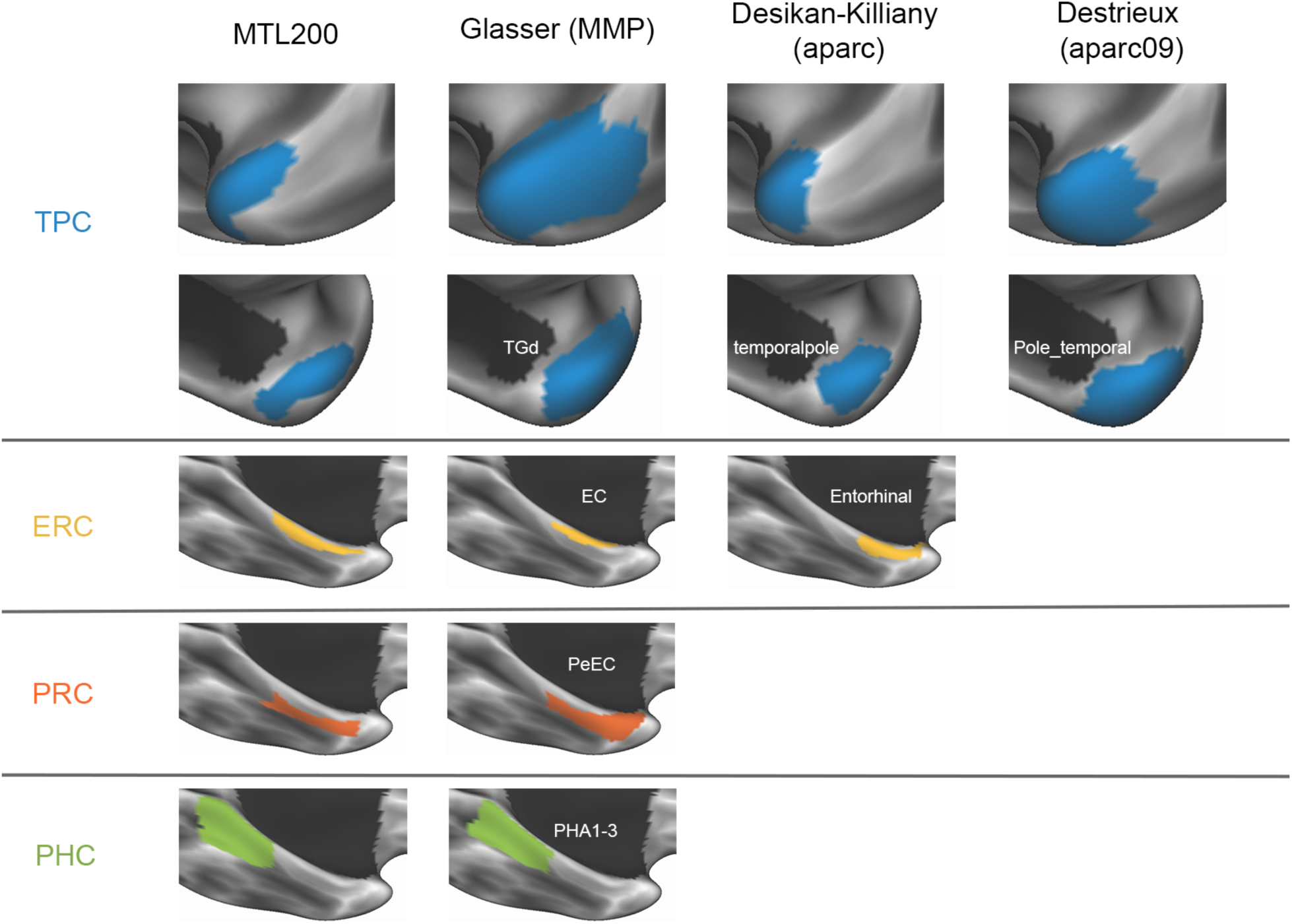
Atlas comparison. Labels for temporopolar cortex (TPC), entorhinal cortex (ERC), perirhinal cortex (PRC), and parahippocampal cortex (PHC) across multiple surface-based atlases: MTL200, Glasser multimodal parcellation, Desikan-Killiany, Destrieux, and MTL200 atlases.

The existing atlases considered here did not have distinct labels for ERC and PRC. MMP includes two regions, labeled entorhinal (EC) and perirhinal/entorhinal (PeEc), while DK includes a single “Entorhinal” region. Since MMP does not differentiate between ERC and PRC per se, we considered the combination of the two areas to compare MMP with MTL200. The combination of MMP’s EC and PeEc corresponded well with the combination of ERC and PRC from MTL200, extending slightly beyond the latter in the anterior direction towards TPC, and the medial direction toward the subiculum. The DK atlas does not include a PRC label, but does include ERC. The boundaries of the “Entorhinal” region in DK did not correspond well with those from MTL200: both anterior and posterior boundaries were positioned anterior in DK relative to MTL200.

Among prior parcellations, MMP was the only one containing a label for PHC; DK and DT include a label for the parahippocampal gyrus but not PHC. The MMP includes three PHC subregions labeled PHA1-3, which are combined in our visualization into a single region. Boundaries of PHC were highly consistent across the MTL200 and MMP. Taken together, these results demonstrate that some MTL region labels in prior cortical parcellations correspond well with boundaries based on cytoarchitecture captured by the MTL200, while others do not. This suggests that caution should be applied when using existing surface-based atlases to identify anatomical regions within the MTL.

### 3.4 Functional connectivity

We next applied the MTL200 to assess the functional connectivity of each region by measuring resting-state correlations. We took two complementary approaches: 1) a “classical” approach using a group analysis, assessing the cross-subject average of correlation maps computed from seed regions defined by the group-level discrete segmentation; and 2) a modern approach in which both MTL regions and functional networks were defined within individuals.

Whole-cortex connectivity maps for each seed region are shown in Fig. 4. TPC showed correlations with areas associated with the default mode network, including medial prefrontal cortex, medial parietal cortex, the temporo-parietal junction, and the superior temporal sulcus. ERC showed the strongest correlations with the occipital lobe, including the lateral, medial, and ventral surfaces. ERC also showed correlations with medial parietal cortex, the precentral gyrus, and regions within the middle and inferior frontal gyri. PRC showed correlations with lateral and ventral occipital cortex, the precentral gyrus, the intraparietal sulcus, and regions within the middle and inferior frontal gyri. PHC primarily showed correlations with occipital regions, including the lateral, medial, and ventral surfaces.

**Figure 4:**
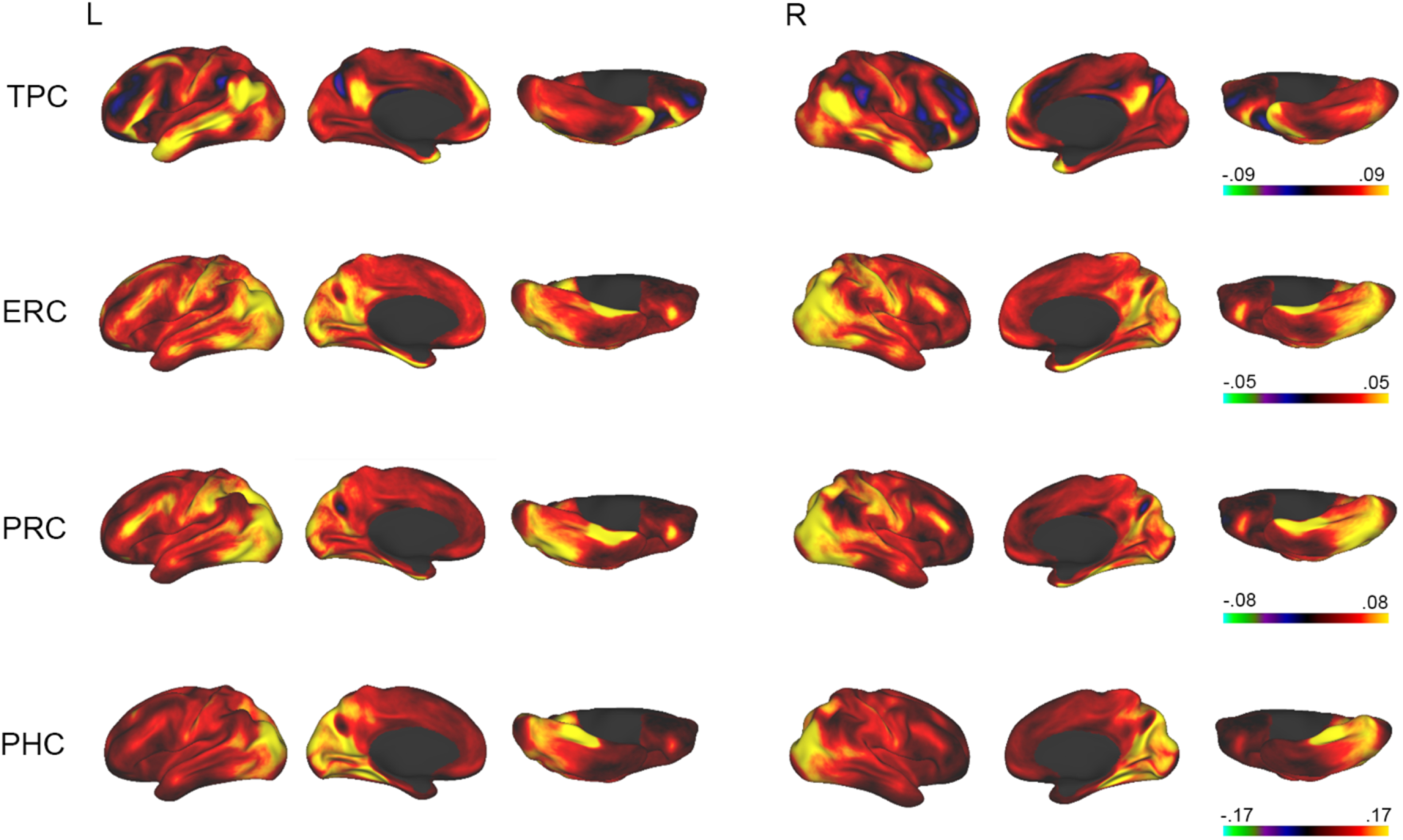
Functional connectivity of MTL regions (group analysis). Maps of correlations with time series from parahippocampal cortex (PHC), perirhinal cortex (PRC), entorhinal cortex (ERC), and temporopolar cortex (TPC). Color map shows correlation strength.

Functional connectivity between individually defined MTL regions and cortex-wide functional networks are shown in Figure 5. TPC had the strongest correlations with default network (DN)-B (DN-B) and the language network (LANG). ERC had a diffuse pattern of correlations across multiple networks, including DN-A, frontoparietal network-A, dorsal attention network (dATN)-A and B, somatomotor networks-A and B, and the auditory network. PRC had the strongest correlations with DN-A. PHC had correlations with DN-A as well as dATN-A and B, the peripheral visual network, and the auditory network. To determine whether patterns of functional connectivity differed significantly across MTL regions, we used a linear mixed model to test for an interaction between seed region and network. The model showed significant main effects of region (*F*[7,23880] = 239.64, *P* < .001) and network (*F*[14,23880] = 16.337, *P* < .001), as well as a significant interaction between region and network (*F*[98,23880] = 56.229, *P* < .001). These results demonstrate that regions within the MTL have dissociable patterns of functional connectivity to distributed networks across cortex.

**Figure 5:**
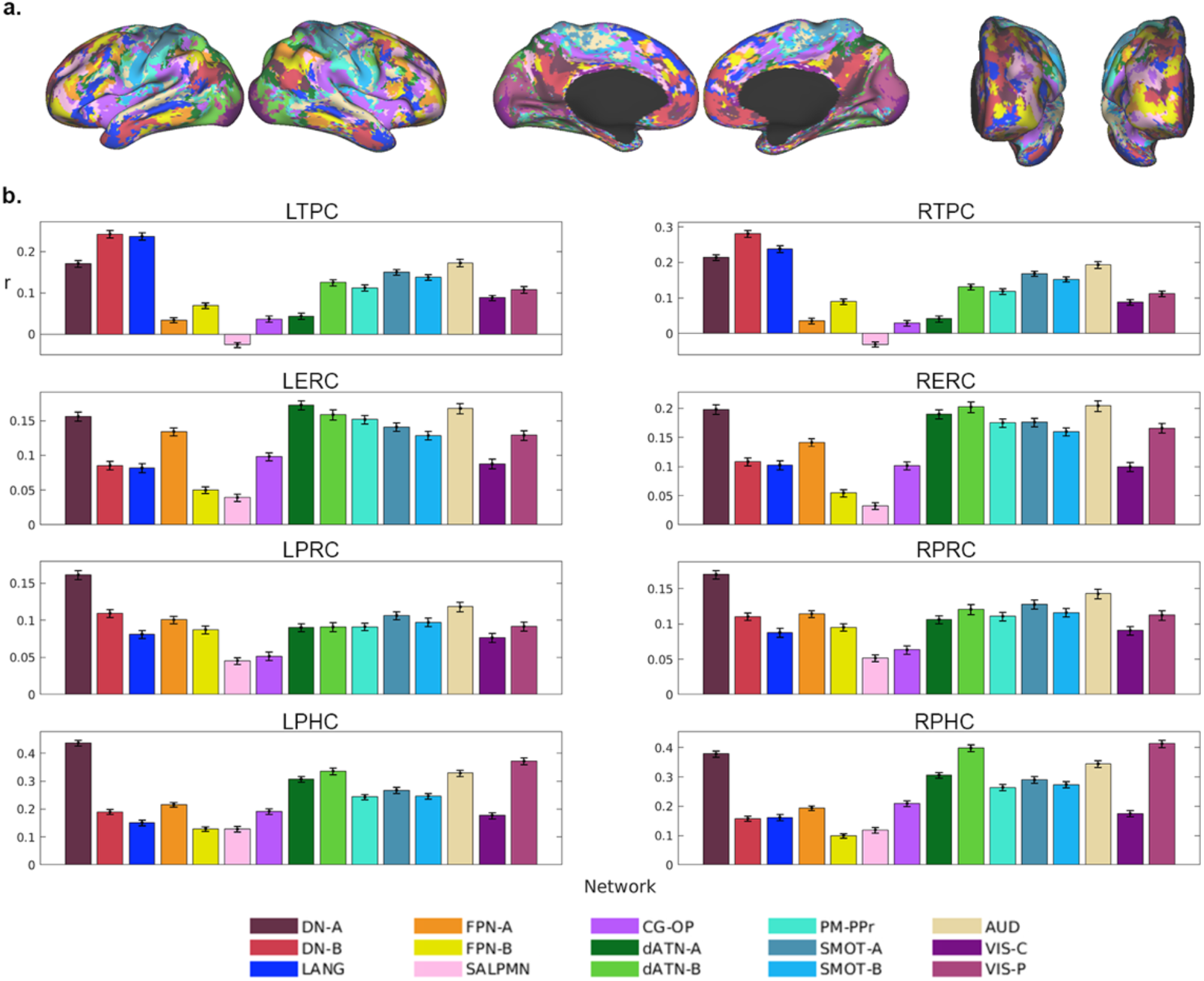
Functional connectivity of MTL regions (individualized network analysis). **a.** 15-network parcellation of cortex within an example participant. **b.** Correlation strength by seed region and network.

## 4. Discussion

We present the MTL200: a surface-based, probabilistic atlas of cortical regions within the MTL, derived from hand drawings in 200 human participants based on gross anatomical features. The atlas is publicly available as a data resource to the neuroimaging community, in order to facilitate the anatomical localization of MTL responses in surface-based analysis of structural and functional MRI data. With a split-half analysis, we show that the atlas is derived from a sufficiently large sample to yield highly reliable probability maps for each region. As an application of the atlas, we assess the functional connectivity of each MTL region based on resting-state correlations, demonstrating dissociable connectivity patterns across areas.

Prior studies providing atlases and segmentation protocols for cytoarchitectonically distinguished MTL regions have taken a volumetric approach (Bouyeure et al., 2018; Yushkevich, Pluta, et al., 2015). Complementing this work, we provide an atlas defined over the cortical surface, expressed in commonly used template spaces fsLR and fsaverage. This atlas will facilitate studies taking the increasingly widespread approach of analyzing fMRI data using cortical surface reconstructions (Bookheimer et al., 2019; Glasser et al., 2013; Hagler Jr et al., 2019; Makropoulos et al., 2018; Somerville et al., 2018; Van Essen et al., 2012). Intersubject registration using surface-based approaches has been found to provide improved alignment of sulci and gyri relative to traditional volumetric approaches (Coalson et al., 2018), which may improve the accuracy of predictions of the location of MTL regions from a surface-based atlas.

As a probabilistic atlas, the MTL200 provides information about both the MTL region likely to exist at a given surface coordinate, and the level of uncertainty in this prediction. We emphasize that this uncertainty results from variation across participants in 1) the location of gross anatomical features; and 2) how these features are registered to fsLR space. Because the atlas was drawn based on gross anatomical rather than cytoarchitectonic features, it does not incorporate variation in the location of cytoarchitectonic boundaries across participants, or variation in the segmentation protocols used to map cytoarchitectonic to anatomical landmarks. Across the literature, somewhat different protocols have been applied to segment the MTL, and harmonizing these different approaches is an active area of research (Wuestefeld et al., 2024; Yushkevich, Amaral, et al., 2015). The current atlas should therefore be interpreted as a description of anatomical variation in areal boundaries determined by a specific segmentation protocol, which was in turn determined based on a cytoarchitectonic analysis (Insausti et al., 1998).

Unlike most prior anatomical atlases of the MTL, the current work includes TPC, a primate-specific region that has been less thoroughly studied than other MTL subregions. To our knowledge, the only existing probabilistic atlas of TPC based on cytoarchitectonic boundaries comes from a volumetric study using a youth sample (Bouyeure et al., 2018). The MTL200 therefore offers a valuable tool for determining whether functional responses in fMRI studies fall within the bounds of TPC. Many prior studies have used Brodmann area boundaries (BA38) to determine whether to characterize a response as falling within the temporal pole (e.g. Ardila et al., 2014; Von Der Heide et al., 2013). However, accounts of the borders of TPC have evolved over time: early maps from Brodmann and von Economo describe an area 38 or TG more extensive than modern accounts, which describe a region restricted to the anterior tip and anteromedial surface of temporal cortex (Blaizot et al., 2010; Ding et al., 2009; Insausti, 2013). The use of a probabilistic atlas grounded in data from recent neuroanatomical work may improve localization accuracy in future fMRI studies.

We also investigate how regions of the MTL200 relate to several widely used surface-based discrete segmentations, based on gross anatomy (DK and DT; Desikan et al., 2006; Destrieux et al., 2010) or connectivity (MMP; Glasser, Coalson, et al., 2016). We found that the temporal pole region from DT and MMP extended farther posterior on the lateral surface of the temporal pole than the bounds of TPC. Given that MMP was derived using patterns of whole-brain anatomical and functional connectivity, this suggests that TPC in humans may have similar connectivity with adjacent regions of far-anterior STS and MTG, consistent with prior reports associating both TPC and anterior STS/MTG with the default mode network (Girn et al., 2024). By contrast, areal boundaries of PHC and ERC/PRC corresponded well across MMP and MTL200, suggesting a correspondence between cytoarchitectonic and connectivity-based boundaries for these areas.

As an initial application of our atlas, we assessed the resting-state functional connectivity of each subregion. PHC showed the strongest correlations with DN-A and VIS-P. These results are consistent with a substantial literature on connectivity between the parahippocampal gyrus and the default mode network (e.g. Greicius et al., 2003; Kahn et al., 2008; Vincent et al., 2007), including recent work associated the parahippocampal gyrus specifically with DN-A and not DN-B (Braga & Buckner, 2017). Correlations with VIS-P are also consistent with work finding dissociable patterns of functional connectivity within the parahippocampal gyrus, with more posterior regions associated with visual areas with a peripheral retinotopic bias (Baldassano et al., 2013; Baldassano et al., 2016; Steel et al., 2021). The present results indicate that these two patterns of functional connectivity exist within the bounds of PHC and not just the broader region of the parahippocampal gyrus.

ERC and PRC showed relatively weak correlation strengths, with less clear differentiation across networks. This pattern may be explained by two methodological factors. First, ERC and PRC have low signal quality due to magnetic susceptibility artifacts affecting fMRI data from the ventral anterior temporal lobes. Second, recent work has found that distinct subregions of ERC and PRC are functionally coupled to separate networks (Reznik et al., 2024), which the current analysis would have combined. Nevertheless, group-level analyses showed correlations with occipital areas, the intraparietal sulcus, and lateral frontal cortex, similar to patterns previously observed for PRC (Reznik et al., 2023). Future work may use the anatomical boundaries provided by the MTL200 as guidelines to identify subregions for further study.

TPC showed the most clearly distinguished connectivity profile, showing correlations with networks DN-B and LANG—distinct but spatially adjacent systems implicated in social cognition and language, respectively (Braga et al., 2020; DiNicola et al., 2020; Paunov et al., 2017). These results are consistent with prior reports associating medial regions of the temporal pole with the default mode network (Deen et al., 2024; Girn et al., 2024; Pascual et al., 2015), as well as work showing anatomical connections with medial prefrontal cortex in the macaque (Moran et al., 1987). Functional connectivity between TPC and LANG has not been emphasized in prior studies. Given that our approach averages signal across TPC, future work should investigate whether distinct subregions of TPC are preferentially connected to DN-B compared to LANG.

In summary, the current work provides a surface-based, probabilistic atlas of cortical regions within the MTL as a data resource. We advance prior work by using a large sample size, generating probability maps on the cortical surface, and including TPC. We hope that this atlas will facilitate neuroanatomically precise study of this important region at the interface of neocortex and the hippocampal formation.

## Acknowledgements

This project was supported by a grant from the Louisiana Board of Regents (LEQSF[2024-27]-RD-A-25 to B.D.). Data were provided by the Human Connectome Project, WU-Minn Consortium (Principal Investigators: David Van Essen and Kamil Ugurbil; 1U54MH091657) funded by the 16 NIH Institutes and Centers that support the NIH Blueprint for Neuroscience Research, and by the McDonnell Center for Systems Neuroscience at Washington University.

## Data/code availability

The atlas and code used for its generation are available at https://github.com/SocialMemoryLab/MTL200. Data used to generate the atlas come from the Human Connectome Project 1200 subjects data release (https://www.humanconnectome.org/study/hcp-young-adult/document/1200-subjects-data-release).

## Author contributions

B.D. conceptualized the project and provided supervision. B.D., A.A., A.S. conducted project administration and data curation. I.F., S.G., A.N., J.P., and K.T. generated hand drawings. I.F. and B.D. conducted data analysis and wrote the manuscript.

