## Supplementary Information for "The MTL200: a surface-based, probabilistic atlas of the medial temporal lobe"

Supplementary Information for Faul et al. “The MTL200: a surface-based, probabilistic atlas of the medial temporal lobe.”

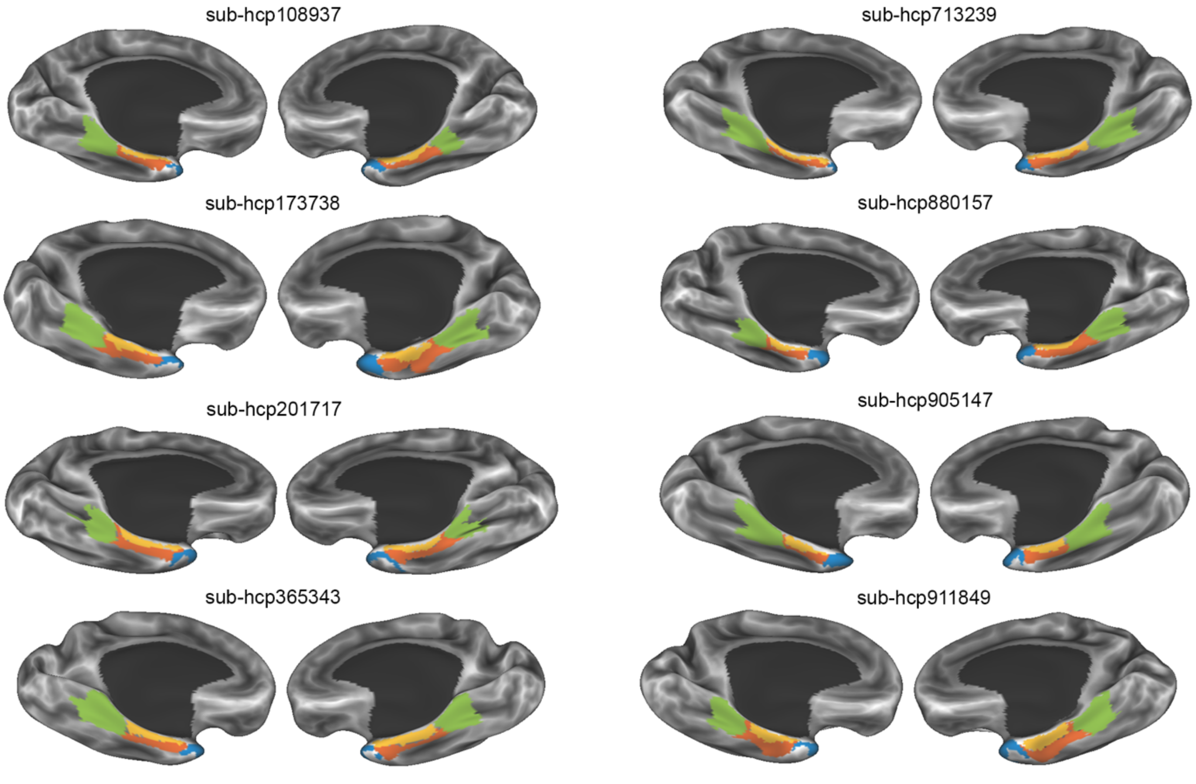


**Fig S1.** Parcellations on the cortical surface in eight example participants (medial view).

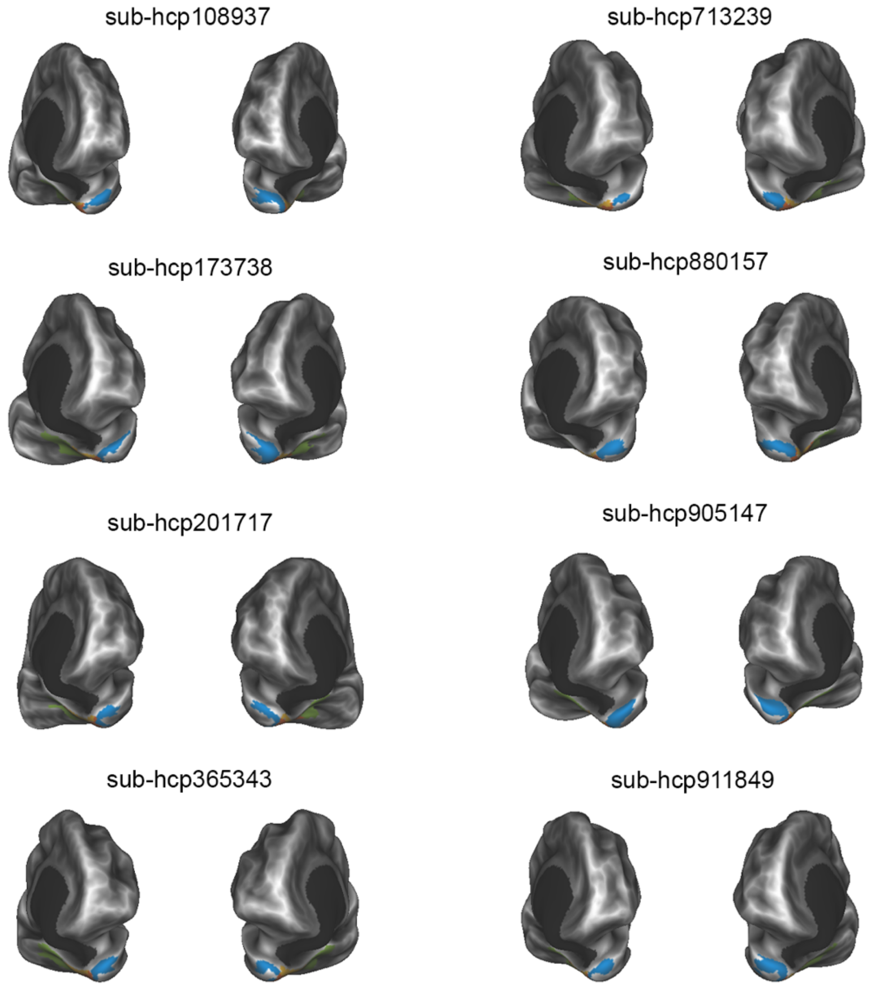


**Fig S2.** Parcellations on the cortical surface in eight example participants (frontal view).

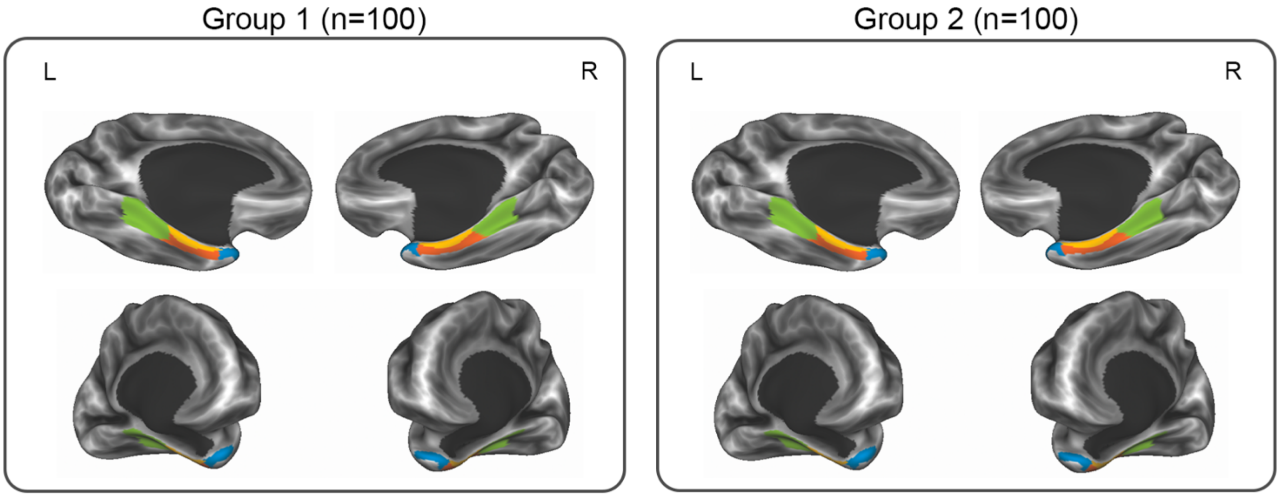


**Fig S3.** Split-half analysis demonstrates high reliability in the discrete segmentation generated across two partitions of participants.

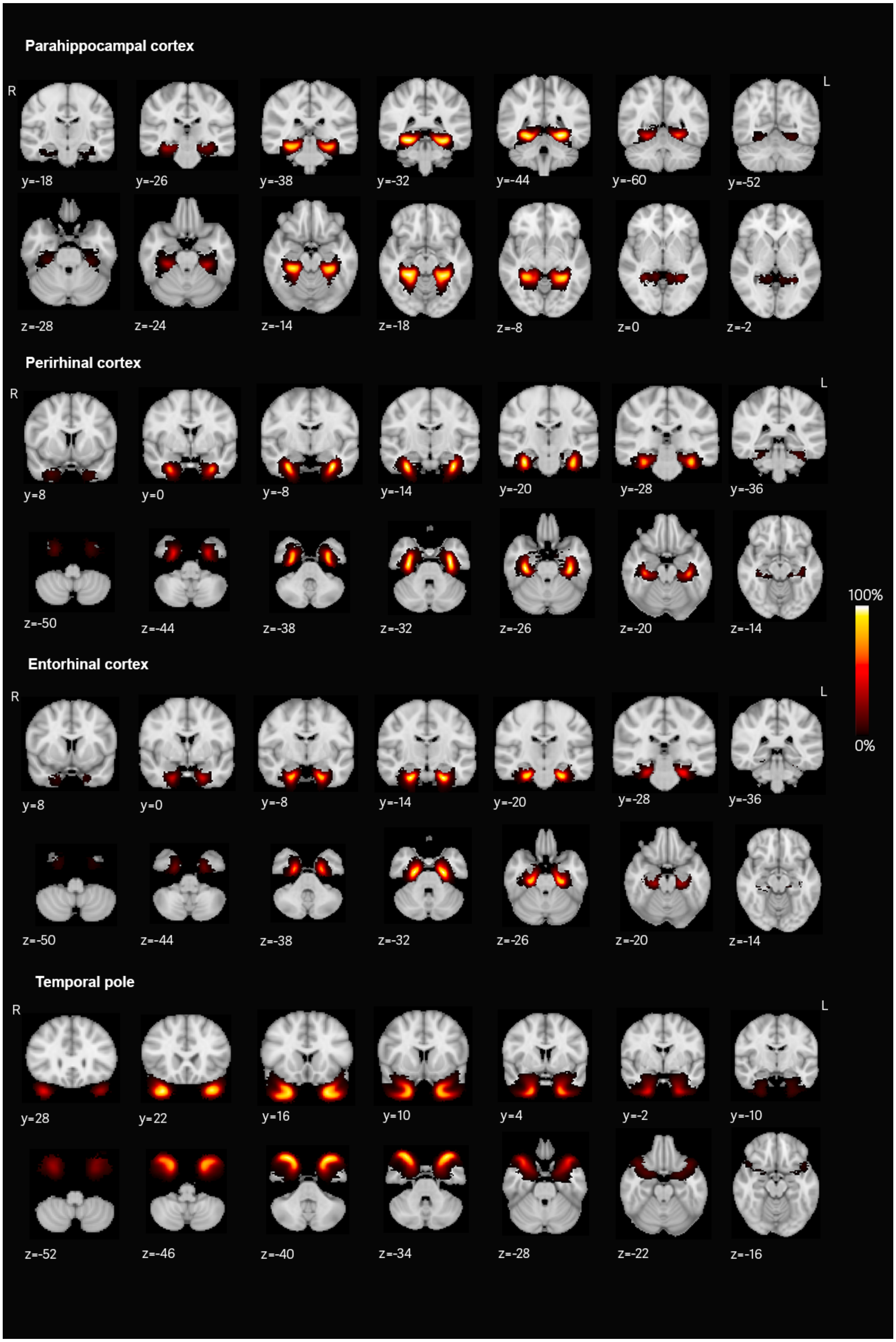

Parahippocampal cortex
R
L
y=-18
y=-26
y=-38
y=-32
y=-44
y=-60
y=-52
z=-28
z=-24
z=-14
z=-18
z=-8
z=0
z=-2
Perirhinal cortex
R
L
y=8
y=0
y=-8
y=-14
y=-20
y=-28
y=-36
z=-50
z=-44
z=-38
z=-32
z=-26
z=-20
z=-14
100%
0%
Entorhinal cortex
R
L
y=8
y=0
y=-8
y=-14
y=-20
y=-28
y=-36
z=-50
z=-44
z=-38
z=-32
z=-26
z=-20
z=-14
Temporal pole
R
L
y=28
y=22
y=16
y=10
y=4
y=-2
y=-10
z=-52
z=-46
z=-40
z=-34
z=-28
z=-22
z=-16

**Fig S4.** Volumetric implementation of the probabilistic atlas in MNI space.
